# GPR158 in pyramidal neurons mediates social novelty behavior via modulating synaptic transmission in mice

**DOI:** 10.1101/2023.04.09.536148

**Authors:** Shoupeng Wei, Jinlong Chang, Jian Jiang, Jin Zhang, Dilong Wang, Yiming Li, Shuwen Chang, Huiliang Li, Ningning Li

## Abstract

Social novelty impairment is a hallmark feature of autism spectrum disorder associated with synaptic dysfunction. While G-protein coupled receptor 158 (GPR158) has been shown to be essential for synaptic neurotransmission, its role in modulating social novelty remains unknown. Here, we investigated the impact of GPR158 on social behavior in mice and observed that both constitutive and cell/tissue-specific knockout of *Gpr158* in pyramidal neurons or the medial prefrontal cortex (mPFC) result in impaired novelty preference, but not sociability. Notably, we found a significant decline in excitatory synaptic transmission and glutamate vesicles in the mPFC synapses of global *Gpr158* knockouts. Mechanistically, we identified that constitutive loss of *Gpr158* led to suppressed Vglut1 distribution, possibly resulting from altered expression of vesicular V-ATPases and SNAREs by *Gpr158* ablation in pyramidal neurons. Our findings suggest that GPR158 in pyramidal neurons specifically modulates social novelty and may be a potential therapeutic target for treating social disorders.

**Graphic Abstract:** 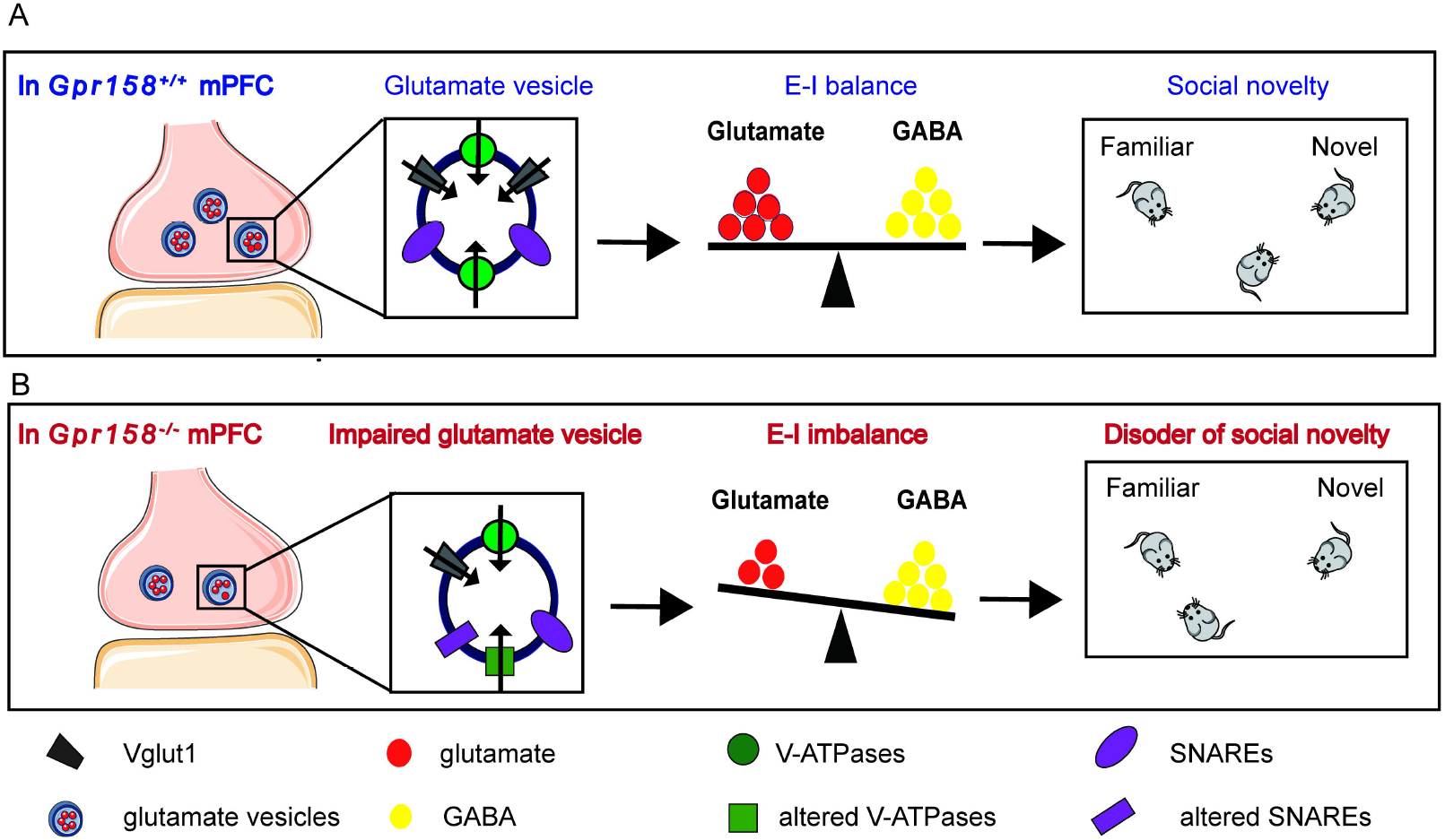

**Highlights:**

1. Knockout of *Gpr158* causes social novelty deficit in mice.
2. Knockout of *Gpr158* in pyramidal neurons leads to disrupted synaptic transmission and an E-I imbalance in the mPFC.
3. Disturbed E-I homeostasis in the mPFC is likely due to reduced density of glutamate vesicles caused by *Gpr158* knockout.

## Introduction

Social interaction is critical to recognize novel stimuli and learn new knowledge, an essential step for being integrated into social life and survival for mammals^1–4^. Behaviorally, social interaction is defined as a phenotype of communication between conspecifics, including two aspects: (1) sociability, a tendency to seek social communication; (2) social novelty, a propensity to communicate with an unfamiliar conspecific^3, 5, 6^. In clinic, two core symptoms, namely impaired social ability and stereotyped behaviors, are adopted to characterize autism spectrum disorder (ASD) as a neurodevelopmental disease^2, 7–9^. ASD accounts for a prevalence rate of approximately 1% worldwide, suggesting that a large population are currently suffering from social disorder^8, 10^. However, the neurobiological mechanism underlying social disorder remains understudied.

Impaired excitation-inhibition (E-I) homeostasis within the medial prefrontal cortex (mPFC) has been demonstrated to be one of the mechanisms underlying the abnormal phenotype of social interaction in rodents^2, 3, 5, 11, 12^. It has been reported that a reduced glutamate/GABA ratio within the PFC and hippocampus is correlated to social deficit in *Cntnap2*^−/−^ mice, a model of ASD. In particular, the mPFC is a key brain area for social behavior^5, 11^. For example, *Cao* et al found that the disrupted gamma oscillations caused by dysfunctions of cortical parvalbumin (PV)-expressing basket cells of the mPFC lead to social novelty deficit in mice^3^. In ASD patients, E-I imbalance and synaptic deficits are frequently accompanied with disrupted gamma bands in the PFC^10, 13, 14^. The extracellular level of neurotransmitter is determined by velocity and quantity of neurotransmitter release and reuptake^15–20^. Many proteins located in the vesicular membrane play a role in this physiological process, e.g., vesicular glutamate transporters (Vglut1) for transporting glutamate, vacuolar ATPases (V-ATPases) for providing energy and a proton electrochemical gradient, and vesicle-associated V-soluble N-ethylmaleimide-sensitive factor attachment protein receptors (SNAREs) for directing the vesicles and facilitating fusion to the presynaptic membrane^15–20^. In short, the vesicular proteins are essential for maintaining E-I homeostasis.

G-protein coupled receptor 158 (GPR158) is a protein expressed by the gene *Gpr158* located in Chromosome 10 of *Homo sapiens,* and it is involved in synaptic function and highly expressed in neuronal cells of the mPFC and hippocampus^21–23^. Our results have shown that GPR158 is mainly distributed in cortical cortex and only expressed in neurons within the central nervous system (CNS)^24^. Loss of *Gpr158* in the mPFC causes an anti-depressant effect in mice, while GPR158 participates in presynaptic maturation of dendritic spines and synaptic formation in hippocampus and cortex^21, 25–27^. Regulator of G Protein Signaling 7 (RGS7) is recruited by GPR158 to the plasma membrane and regulates the generation of intracellular adenylate cyclase (AC) and cyclic adenosine monophosphate (cAMP) in the state of stress^21, 26, 28^.

In this study, we investigated the role of GPR158 in social novelty by knocking out *Gpr158* in mice. We found that abnormal structures and dysfunction of glutamatergic transmission underlie social novelty deficit in the *Gpr158* knockouts. In addition, Vglut1 distribution in the mPFC was reduced when *Gpr158* was globally knocked out, while expression of V-ATPases and SNAREs was interrupted in mice with conditional deletion of *Gpr158* in pyramidal neurons. In short, we revealed the mechanism of GPR158 modulating social novelty behaviors, providing an in-depth understanding of the neurobiological substrates underlying social disorder.

## Materials and Methods

### Animals

All mice adopted in this study were on a C57Bl/6J background and housed in a pathogen-free SPFII facility with constant temperature (23±1 ℃) and controlled humidity (50±10%) on a 12h/12h light/dark cycle (lights on at 07:00) in the Center of Laboratory Animal Science, Southern University of Science and Technology (SUSTec), China. Male mice were 8-10 weeks old, weighing approximately 20 g, at the beginning of behavioral tests. Mice were maintained in a group of 4 to 5 in a clear plastic cage, and had free access to drinking water and standard food. All experiments were conducted according to the NIH Guide for the Care and Use of Laboratory Animals (NIH publications no. 80-23, revised 1996), and the procedures were approved by the Animal Care Committee at the SUSTec, China.

In this study, *Gpr158^-/-^* (*Gpr158^tm^*^1^(KOMP)*^Vlcg^*) and *Gpr158^fl/fl^* mice were imported from Columbia University (NY, USA), while *CamK2A-Cre*, *Vgat-Cre* and *Gpr158-e(3×Flag-TeV-HA-T2A-tdTomato)1* mouse lines were bought from Shanghai Model Organisms Company, China^21, 25^. *Gpr158* is located in the chromosome 2 of *musculus* and harbors 11 exons. The primers used to genotype the transgenic mice are listed in Supplementary Table 1. Mouse DNA was extracted from mouse tails for polymerase chain reaction (PCR) with Quick Genotyping Assay Kits for Mouse Tail bought from Beyotime Biotechnology (Shanghai, China). The first 2 exons of *Gpr158* were replaced with a *LacZ* cassette in *Gpr158^-/-^* mice. *CamK2A-Cre;Gpr158^fl/fl^* and *Vgat-Cre;Gpr158^fl/fl^* mice were, respectively, obtained by crossing *CamK2A-Cre* and *Vgat-Cre* mice with *Gpr158^fl/fl^* mice, where the exon 11 was flanked by the *loxP* sites and deleted by Cre-induced recombination^29^. The promoters of *CamK2A* encoding calcium/calmodulin-dependent protein kinase II alpha (CamK2A) and *Vgat* encoding vesicular gamma-aminobutyric acid (GABA) transporter (Vgat), were used to control *Cre* expression^9,30–32^. *Gpr158-e(3×Flag-TeV-HA-T2A-tdTomato)1* mouse strain was established to indicate the expression of *Gpr158,* co-expressed with the Tag genes *Flag, HA* and *tdTomato* in this study.

### Microinjection

Mice were anesthetized with 1.5-1.8% isoflurane and fixed on a stereotaxic frame (RWD Life Science Co., Ltd, Shenzhen, China). Recombinant adeno-associated virus (rAAV) was bilaterally injected into the mPFC (from the bregma: anterior-posterior +1.8 mm, medial-lateral ±0.4 mm, dorsal-ventral -2.6 mm) with a 10µl Hamilton microsyringe. The volume of each rAAV microinjection was set at 0.5 µl and delivered at a speed of 1.5 nl/s to minimize diffusion. The injector was kept for 10 min after the injection to prevent fluid back flow, and then slowly withdrawn. The incision was sutured with sterile string and closed with tissue glue. For silencing *Gpr158* expression in the mPFC of wild-type (WT) mice, *rAAV-U6-shRNA-Gpr158-CMV-EGFP-sv400pA* virus (*shRNA-Gpr158*) and *rAAV-CMV-EGFP-sv400pA* virus (*shRNA-Scramble*) with high VG titer (≥ 1 × 10^13^ vg/ml) were used. For knocking out *Gpr158* within pyramidal neurons of *Gpr158^fl/fl^* mice, *rAAV-CamK2A-Cre* virus (*rAAV-CamK2A-Cre*) and *rAAV-CamK2A-eGFP* virus (*rAAV-Scramble*) with high VG titer (≥1 × 10^13^ vg/ml) were used. The mice were given 3 weeks after the virus injection for viral expression, and the knockdown efficiency was assessed using quantitative polymerase chain reaction (qPCR) assay.

### Three-chamber social interaction test (TCT)

TCT was carried out on age-matched male mice during the daytime as described somewhere^3,4^. The apparatus (60 cm long × 40 cm wide × 25 cm high) consisted of 3 chambers with each side chamber of 20 cm long. The paradigm consisted of 3 phases:

(1) habituation. One male mouse was gently placed in the center chamber for 5 min, with both of the entrances blocked by guillotine doors. (2) sociability. One stranger male mouse (S1) was put inside a wire cage in the central area of the chamber, while another identical empty wire cage (E) was put at the symmetric position in another side chamber. The guillotine doors were then opened, allowing the habituated experimental mouse to explore both side chambers for 10 min. (3) social novelty. The mouse was guided into the center chamber, and another stranger mouse (S2) was put inside the previous empty cage. The mouse was given another 10 min to explore the whole apparatus. In this test, S1 and S2 were at the similar size and age compared to the experimental mice. The apparatus was cleaned with 75% ethanol between the sessions. A wire cage to restrain a mouse was at a diameter of 8 cm, and a circle at the diameter of 14 cm shared the same center was set as the interaction area. Behaviors were recorded and analyzed with Noldus EthoVision XT10 software (Wageningen, The Netherlands), and the interaction time spent with S1 *versus* E in the second phase was analyzed to indicate their sociability, and with S1 *versus* S2 in the third phase to indicate their social novelty preference.

### Marble burying test (MBT)

MBT is performed to assess stereotyped behaviors of rodents^6,21, 33^. Briefly, mice were habituated for and tested in the transparent Plexiglas cages (37 cm long × 14 cm wide × 12.5 cm high) with 5cm thick bedding material. After 30min habituation, mice were placed in another cage with 20 glass marbles (at the diameter of 1.5 cm) placed on the bedding at 4 × 5 rows, freely exploring the cage for 10 min. The number of buried marbles were counted after the test. Marbles with 75% of the surface covered by the bedding were defined as being buried.

### Novel object recognition test (NORT)

NORT is carried out to assess the ability to distinguish the familiar and novel objects and to evaluate learning ability and social memory in rodents^21, 34^. The NORT paradigm was composed of 2 phases: (1) familiarization. A mouse was put into the open field box (40 long× 40 cm wide× 45 cm high) and allowed free exploring the area for 5min where two identical objects (#A and #B) were placed at a distance of 10 cm from each other. (2) test. The mouse was taken out and put in the home cage, and then re-exposed to the area after 2 h, where one of two objects #B was replaced by a novel object #C for 5 min. The round zone at a diameter of 3 cm to the object was set as the recognition area, and the time spent in the recognition area was recorded by with Noldus EthoVision XT10 software (The Netherlands). The apparatus was cleaned with 75% ethanol between sessions. Recognition Index (RI) = T_novel_/(T_novel_ + T_familiar_). T_novel_ and T_familiar_ representing the time spent in the recognition area of the novel and familiar object, respectively.

### Enzyme-linked immunosorbent assay (ELISA)

ELISA was performed according to the manufacturer’s instructions^31, 32^. Dissected brain tissues and serum were stored in -80 ℃ for use. To lyse tissues, ice cold 1× Phosphate Buffered Saline (PBS) was added based on tissue weight and homogenized on ice. The supernatant was collected after the samples were centrifuged at ∼13400 g/min at 4 ℃ for 30 min. The levels of glutamate, GABA and dopamine were measured by a sandwich ELISA using 100µl serum or the supernatants. Optical density was measured by 450nm light with an ELISA microplate reader (Thermo Fisher Scientific, MA, USA).

### Immunofluorescent (IF) staining

The protocol for IF staining previously published was adopted in this study^1,3^. Mice were transcardially flushed with 1× PBS and then perfused with 4% paraformaldehyde (PFA) after being anesthetized with isoflurane. Next, the brains were post-fixed in 4% PFA at 4 ℃ overnight, cryoprotected with 20% and 30% sucrose for 12-24 h, respectively. They were embedded in Tissue-Tek OCT (Sakura Seiki, Nagano, Japan) and stored at −80 ℃ for cutting. Brains were sectioned at 30 μm. Targeted brain slices were first rinsed with 1× PBS 3 times (10 min/each) and blocked with a blocking solution, 10% goat serum and 0.2% Triton X-100 in PBS, for 1 h at room temperature. Brain slices were incubated with the primary antibodies prepared in a blocking solution at 4 ℃ overnight. On the next day, slices were rinsed with 1× PBS 3 times and incubated with the secondary antibodies prepared in a blocking solution for 90 min. Then, the slices were washed with 1× PBS for 10 min, and then incubated with 1 μg/ml DAPI for 10 min. Brain slices were washed with 1× PBS 3 times before wet-mounted on a slide and coverslipped with the fluoroshield mounting medium. Slides were imaged with a ZEISS LSM880 confocal microscope (Jena, Germany). Tissues for immunostaining c-Fos proteins expressed by immediate early genes, were harvested 90 min after the social novelty test, while the tissues for other immunostaining tests were collected immediately following the test.

### Transmission electron microscope (TEM) photography

The protocol for TEM was modified from the previous publications^35, 36^. Brains of anesthetized mice were quickly removed and put in pre-cooling PBS containing 2.5% glutaraldehyde. The mPFC tissues were collected and kept in fresh TEM fixative (Servicebio, Wuhan, China) at 4°C overnight. Tissues were washed and fixed with 1% OsO4 in 0.1 M Phosphate-Buffered Saline (PBS) for 2 h at room temperature in the dark room followed by being washed with 0.1M PBS (15 min each) for 6 times to get rid of OsO4. Next, the tissues were dehydrated by a series of ethanol solutions (30%, 50%, 70%, 80%, 95% and 100%) 2 times (20 min each) followed by acetone 2 times (5 min each) at room temperature. The tissues were embedded in resin to polymerize at 65 °C for 48 h. Then, the tissues were sectioned at 60-80 nm with a Leica EM UC7 ultramicrotome (Wetzlar, Germany). The sections were mounted onto the formvar-coated 150-mesh copper grids for photography by a Tecnai G2 Spirit TEM (Thermo Fisher Scientific, MA, USA). Synaptic substructures were observed at the magnification power of 26,400 using TEM.

### Golgi-Cox staining

Hito Golgi-Cox Kit (Hitobiotec Corp., Wilmington, DE, USA) was used to reveal the structure and density of dendritic spines in the mPFC^37–39^. Mice were killed, and brains were harvested and rinsed in cold 1× PBS for 2-3 seconds. The brains were immersed into a mixture of solutions 1 and 2 for 24 h, replaced with the fresh mixture and stored for 2 weeks in the dark at room temperature. The brains were transferred into solution 3, stored for 72 h and then cut into the slices of 150 µm thick with a Leica CM3050S cryostat (Wetzlar, Germany). Slices were mounted on gelatin-coated slides, stained with a mixture of solution 4, 5 and 6, dehydrated with 50%, 75%, 95% and 100% ethanol, cleared by xylene and coversliped with resinous mounting medium. After drying, the slides were taken photos of with a bright field microscopy.

### Electrophysiological recording

Whole-cell clamp patch was used to assess the neuronal physiological properties^3, 6, 25^. Mouse brains were quickly removed and put in a pre-cooling cutting solution with being continuously aerated with a mixture of 95% CO_2_ and 5% O_2_. Brains were coronally sectioned at 300 μm with a Leica VT1200S vibratome (Wetzlar, Germany), and the slices were put into standard recording artificial cerebrospinal fluid (ACSF) at 37 ℃ for 30 min. To record it, a slice was put in a superfusion chamber, secured under a nylon mesh and superfused at 2 ml/min with the ACSF. A multiclamp700B amplifier was used, and data from the neurons would not be used if the series resistance was more than 30 MΩ, or depolarization of the membrane potential was higher than -45 mV at the onset of breaking in, or the baseline was not stable with the alteration larger than 20% during recording. Here, excitatory postsynaptic currents (EPSCs) were recorded in the mPFC layer 2/3 area. The cutting solution was prepared following the format: 2.5 mmol/L KCl, 1.25 mmol/L NaH_2_PO_4_, 26 mmol/L NaHCO_3_, 8 mmol/L MgSO_4_, 0.5 mmol/L GaCl_2_, 10 mmol/L Glucose, 115 mmol/L Choline chloride, 0.1 mmol/L Ascorbic acid, the solution adjusted to the osmotic pressure of 300-305 mOsm/L and the PH of 7.35-7.45. The artificial spinal fluid (ACSF) was prepared following the format: 125 mmol/L NaCl, 2.5 mmol/L KCl, 1.25 mmol/L NaH_2_PO_4_, 26 mmol/L NaHCO_3_, 1.3 mmol/L MgSO_4_, 2.5 mmol/L GaCl_2_, 10 mmol/L Glucose, 2 mmol/L Na-pyruvate, 0.5 mmol/L Ascorbic acid, 3 mmol/L Myo-inositol, the solution adjusted to the osmotic pressure of 300-305 mOsm/L and the PH of 7.35-7.45.

### Western blotting (WB)

WB assay was performed according to the manufacturer’s instructions. Brain tissues were dissected and used to extract the total protein using RIPA lysis buffer (Beyotime, Shanghai, China). The supernatant was collected after the sample was centrifuged at ∼13400 g/min at 4 ℃ for 30 min. Bicinchoninic acid (BCA) method was used to quantify the concentration of total protein. Targeting proteins were separated using sodium dodecyl sulfate-polyacrylamide gel electrophoresis (SDS-PAGE) by electrophoresis and transferred onto a polyvinylidene fluoride (PVDF) film (Merck Millipore, Guangzhou, China). The PVDF membrane carrying proteins was blocked by 5% fat-free milk dissolved in 1× TBST (0.1% Tween20 in Tris-buffered saline) for 1 h at room temperature. Targeted strips were incubated with the primary antibodies dissolved in 5% fat-free milk at 4 ℃ overnight. On the next day, the strips were incubated with HRP-conjugated secondary antibodies after being washed with 1× TBST 3 times (10 min each). Then, 4’,6-diamidino-2-phenylindole (DAPI) was used to stain cell nuclei for 10 min, and the strips were washed with 1× TBST 3 times (10 min each). Immunodetection was carried out with a super ECL detection reagent and detected with the ChemiDoc™ Touch Imaging System (Bio-Rad, Shanghai, China).

### RT-qPCR

Total RNA was extracted from brain tissues, and complementary DNA was obtained by reverse transcription. Real-time quantitative PCR (RT-qPCR) was carried out by SYBR Green detection in a two-step reaction using the Bio-Rad CFX96 Detection System (Applied Bio-systems, USA). Threshold cycle (CT) values were calculated to quantify target gene mRNA, while ΔCT indicated the difference of CTs between the target and reference genes. ΔΔCT represented the difference of ΔCTs between treatment and control group to estimate the expression of target mRNA.

### RNA-sequencing analysis

Total RNA was extracted from the mPFC tissues of *CamK2A-Cre;Gpr158^fl/fl^* mice and the controls (n = 4) at the age of 8 weeks with Tripure Isolation Reagent (Roche, Mannheim, Germany). RNA-sequencing (RNA-seq) was carried out using the Illumina NovaSeq 6000 platform by Novogene Company (Beijing, China). Paired-end clean reads were aligned to the mouse reference genome (Ensemble_GRCm38.90) with TopHat (version 2.0.12), and HTSeq-count (version 0.6.1) was used to quantify mRNA expression by the aligned reads. Differentially expressed genes (DEGs) were picked up for further analysis. The pathways were enriched using the assays of gene ontology (GO) and Kyoto Encyclopedia of Genes and Genomes (KEGG).

### Statistical Analysis

GraphPad Prism8.0 (San Diego, USA) was used to perform statistics and plot figures in this study. One-way or two-way ANOVA was performed to compare the differences among 3 groups or more, followed by LSD *post hoc* tests for multiple comparisons. Unpaired *t* tests were used to compare the difference between 2 groups. Data are presented as mean ± SEM. A significant difference was set as p < 0.05.

## Results

### Knockout of *Gpr158* induced social novelty deficit in mice

To explore the role of GPR158 in social behavior, we employed the global knockouts of *Gpr158* (*Gpr158^-/-^*, KO) to study loss-of-function effect in sociability and social novelty (Figure 1A-C). Mice were subjected to TCT which has been widely used to assess sociability and social novelty in rodents^1, 3, 25^. We found that *Gpr158* KO did not affect social preference to a conspecific over an empty cage (WT: p < 0.01; KO: p<0.01) analyzed with One-way ANOVA, suggesting that *Gpr158* loss did not undermine the phenotype of sociability in mice (Figure 1B). Of significance, KO mice showed the abnormal phenotype of social novelty preference, that is, the duration of social interaction with a novel conspecific *versus* a familiar one was significant shorter than that in WT group (WT: p < 0.01; KO: p = 0.167; Figure 1C). Next, MBT was performed to assess the potential role of GPR158 in modulating autistic-like stereotyped behaviors as one of the core symptoms of ASD, exhibiting that there was no difference in the number of marbles buried between WT and KO mice (t_32_ = -1.048, p = 0.303; Figure 1D). Similarly, NORT revealed that the recognition indexes did not differ between WT and KO mice (t_31_ = -0.04, p = 0.966; Figure 1E), showing that knockout of *Gpr158* did not affect recognitive ability. Thus, we conclude that GPR158 mediates social novelty behavior, but not play a role in sociability, stereotyped behaviors or cognitive ability.

**Figure 1.**
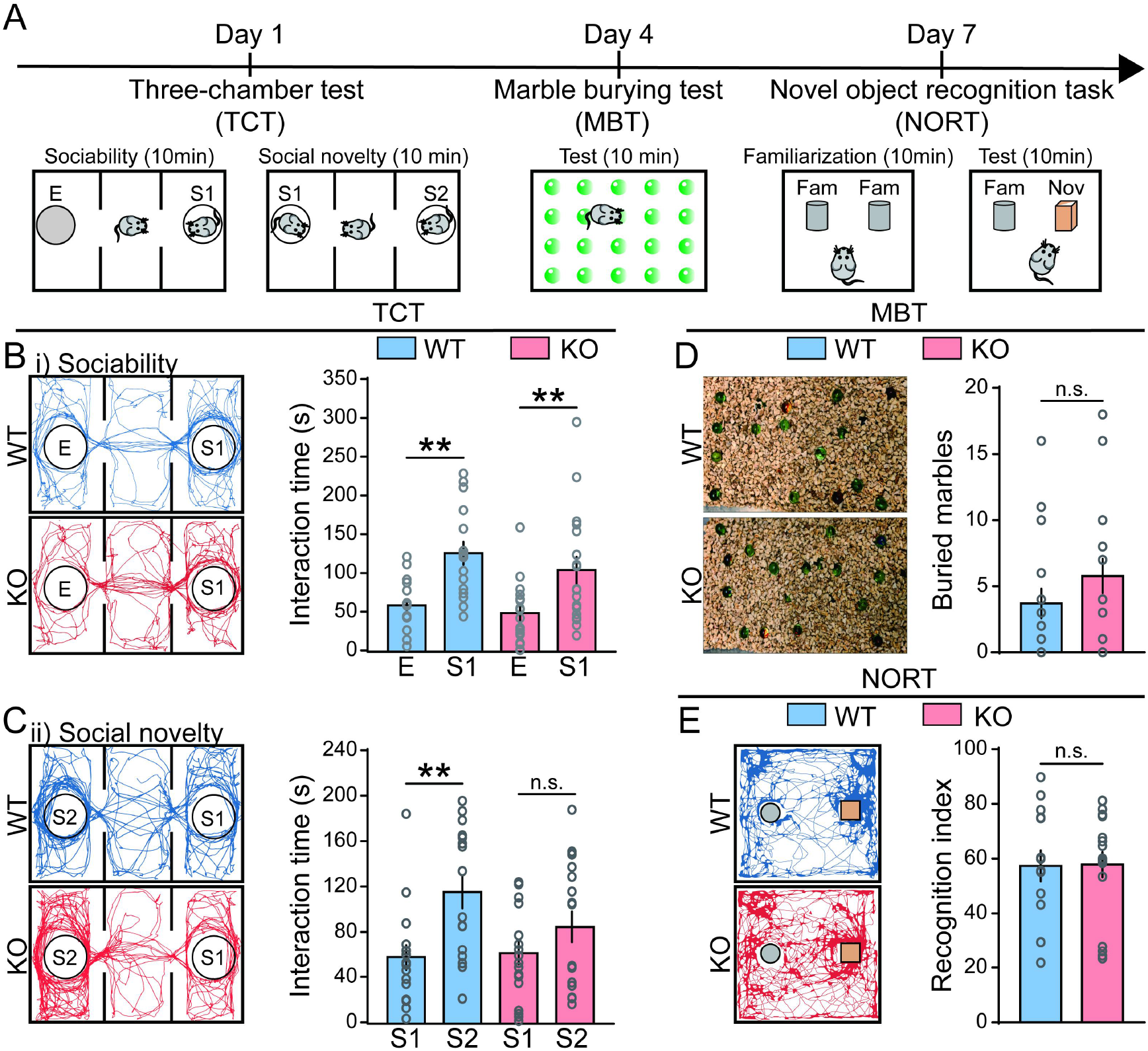
Loss of *Gpr158* induced deficit of social novelty, but not stereotyped behavior or impaired cognitive ability in mice. (A) Procedures of behavioral tests performed in *Gpr158^-/-^* (KO) mice and the WT controls. Three-chamber social interaction Test (TCT), Marble burying test (MBT) and Novel object recognition test (NORT) were subsequently carried out in this study. **(B)** KO mice and the controls both showed significant preference for a stranger (S1) over an empty cage (E) (WT: p < 0.01; KO: p<0.01) in the sociability test of TCT (WT: n=16, KO: n=18), shown in the representative trajectory maps. **(C)** KO mice showed undermined preference for a novel stranger (S2) over a familiar one (S1) (WT: p < 0.01; KO: p = 0.167) compared to that of the control in the social novelty test of TCT, as shown in the representative trajectory maps. **(D)** Representative images of the MBT. Numbers of marbles buried did not differ between KO and WT mice in MBT (t_32_ = -1.048, p = 0.303; WT: n=16; KO: n=18). **(E)** Recognition indexes did not differ between KO mice and WT controls in the NORT (t_31_ = -0.04, p = 0.966; WT: n=14; KO: n=18), shown in the representative images of NORT. F, a familiar object; N, a novel object; WT, wild-type; KO, *Gpr158^-/-^*. n.s., not significant; * p < 0.05, ** p < 0.01. Data were analyzed with One-way ANOVA among 3 groups and the unpaired *t* test between 2 groups. Data are represented as the mean ± SEM.

### Knockout of *Gpr158* reduced the density of c-Fos^+^ neurons in the mPFC

The protein c-Fos expressed by an immediate early gene is used to evaluate neuronal excitation^3, 38–40^. To clarify the neurobiological substrates that mediated social novelty behavior, we searched for the critical target brain area after social behavior task using c-Fos staining. Here, we screened the brain areas including the mPFC, hippocampal CA1, CA3, dental gyrus (DG) and ventromedial hypothalamus (VMH), and found that, by co-labeling c-Fos and NeuN proteins, the reduced density of c-Fos protein in the mPFC and DG of KO mice (mPFC: t_40_ = -4.831, p < 0.01; DG: t_40_ = 2.297, p < 0.01), while the density in hippocampal CA1, CA3 or VMH was not changed (CA1: t_34_ = 1.17, p = 0.096; CA3: t_31_ = -1.674, p = 0.104; VMH: t_14_ = -0.327, p = 0.748) analyzed with unpaired *t* tests, shown in Figure 2A-F. Meanwhile, the density of neurons did not alter in the mPFC of KO mice (t_31_ = 1.271, p = 0.2131), shown in Figure 2G and H. Therefore, the mPFC was chosen as the target area because of the more significant alteration of c-Fos expression triggered by social novelty behaviors. To confirm the *Gpr158* expression in the mPFC, *Gpr158-Tag or Gpr158-e(3×Flag-TeV-HA -T2A-tdTomato)1* mouse was used, suggesting the density of GPR158-expressing cells was 552/mm^2^, and around 50% of the pyramidal neurons in the mPFC expressed GPR158 by fluorescent staining (Supplementary Figure 1). Furthermore, we found that *Gpr158* knockout did not affect the density of pyramidal neurons in the mPFC (t_44_ = 1.125, p = 0.2666; Supplementary Figure 2). To conclude, the neuronal density was not altered in KO mice, showing the reduced density of c-Fos proteins was not attributed to the declining density of neurons.

**Figure 2.**
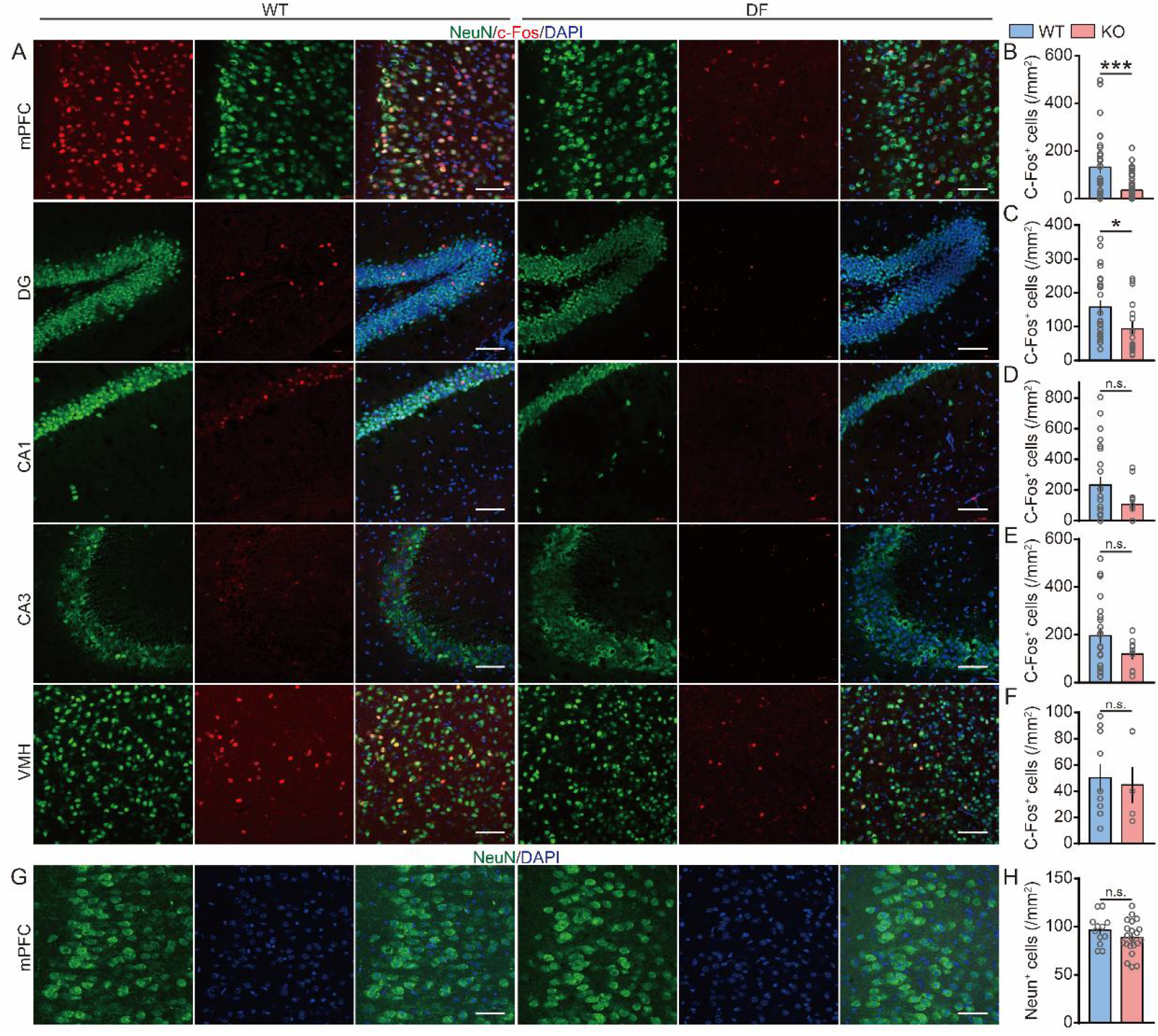
Loss of *Gpr158* reduced density of c-Fos^+^ neurons in the mPFC. Immunofluorescent staining was performed using the tissues collected 90 min after the social novelty test. **(A)** Representative images of c-Fos^+^ cells in the brain areas, namely the mPFC, DG, hippocampal CA1, CA3 and VMH (WT: n=4; KO: n=4). **(B-F)** The density of c-Fos^+^ cells was significantly decreased in the mPFC (t_40_ = -4.831, p < 0.01) and hippocampal DG (t_40_ = 2.297, p < 0.01) of KO mice, but not altered in hippocampal CA1 (t_34_ = 1.17, p = 0.096), CA3 (t_31_ = -1.674, p = 0.104) or VMH area (t_14_ = -0.327, p = 0.748). Scale bar: 20 μm. **(G)** Representative images of neurons (NeuN^+^) in the mPFC of KO and WT mice (WT: n=4; KO: n=4). **(H)** The density of neurons did not change in the mPFC (t_31_ = 1.271, p = 0.2131). Scale bars: 50 μm. mPFC, medial prefrontal cortex; DG, dental gyrus; VMH, ventromedial hypothalamus; WT, wild-type; KO, *Gpr158^-/-^*. n.s., not significant; * p < 0.05, *** p < 0.001. Data were analyzed with the unpaired *t* test. Data are represented as the mean ± SEM.

### Conditional knockout of *Gpr158* within pyramidal neurons induced social novelty deficit

Excitatory CamK2A^+^ pyramidal neurons and inhibitory PV^+^ neurons are two dominant neuronal types in the mPFC, as CamK2A is mainly distributed in excitatory pyramidal neurons, and Vgat is mainly expressed in inhibitory GABAergic and glycinergic neurons^41, 42^. To investigate the subtype of neurons in mediating social novelty behavior, we employed two transgenic mouse lines *CamK2A-Cre;Gpr158^fl/fl^* and *Vgat-Cre;Gpr158^fl/fl^* with *Gpr158* selectively deleted from CamK2A^+^ and Vgat^+^ neurons, respectively. In TCT, we found that *CamK2A-Cre;Gpr158^fl/fl^* (CaKO) mice showed a trend of altered social novelty (CON: p = 0.076; CaKO: p = 0.569), while there was no difference in the phenotype of sociability between CaKO and control mice (CON: p < 0.01; CaKO: p < 0.01), shown in Figure 3A-C. In the contrast, *Vgat-Cre;Gpr158^fl/fl^* (VgKO) mice displayed the similar phenotype of sociability (CON: p < 0.01; CaKO: p = 0.073) and social novelty (CON: p < 0.01; CaKO: p < 0.05), shown in Figure 3D-F. Thus, our data suggest that GPR158 signaling in pyramidal neurons mediates social novelty behavior.

**Figure 3.**
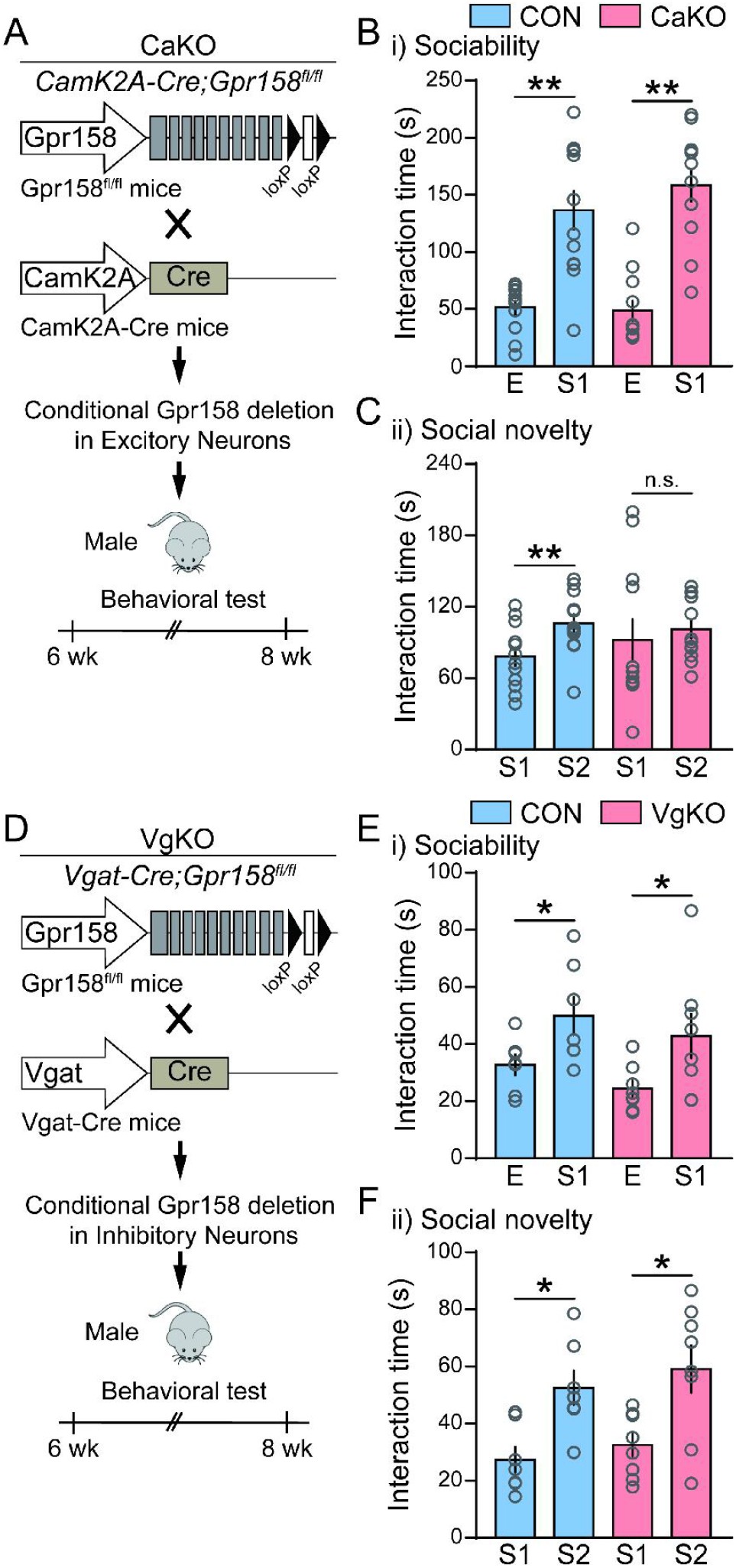
Conditional knockout of *Gpr158* in pyramidal neurons induced social novelty deficit in mice. (A, D) The exon 11 of *Gpr158* gene was selectively deleted by a Cre enzyme in this test. *Gpr158* harbors 11 exons, and the exon 11 was deleted in *CamK2A-Cre;Gpr158^fl/fl^* (CON: n=10; CaKO: n=12) **(A)** and *Vgat-Cre;Gpr158^fl/fl^* mice (CON: n=8; VgKO: n=9) **(D). (B)** CaKO mice and the controls showed social preference to a conspecific over an empty cage in the sociability test compared to the controls (CON: p < 0.01; CaKO: p < 0.01). **(C)** CaKO mice showed altered social novelty preference over a novel stranger (CON: p = 0.076; CaKO: p = 0.569). **(E)** VgKO mice did not show different sociability behavior (CON: p < 0.01; CaKO: p = 0.073). **(F)** VgKO mice did not show different social novelty preference over a novel stranger compared to the controls (CON: p < 0.01; CaKO: p < 0.05). E, an empty cage; S1, stranger 1; S2, stranger 2; F, a familiar object; N, a novel object; n.s., not significant; * p < 0.05, ** p < 0.01. Data were analyzed with One-way ANOVA among 3 groups and the unpaired *t* test between 2 groups. Data are represented as the mean ± SEM.

### Silencing *Gpr158* expression in the mPFC or knocking out *Gpr158* from pyramidal neurons induced social novelty deficit

While using genetic deletion of *Gpr158*, we cannot rule out the effect of GPR158 from other brain regions on social novelty behavior. We further used the virus-based methods to silence *Gpr158* expression in the mPFC and to knock out *Gpr158* from pyramidal neurons, respectively, to validate the specific role of Gpr158 in pyramidal cell in the mPFC. Firstly, the virus *rAAV-U6-shRNA-Gpr158-CMV-EGFP-sv400pA* (*shRNA-Gpr158*) was bilaterally injected into the mPFC of WT mice to silence *Gpr158* expression and *rAAV-CMV-EGFP-sv400pA* (*shRNA-Scramble*) was used as the control. The mice underwent TCT 3 weeks after the virus injection. *ShRNA-Gpr158* significantly inhibited social novelty preference to the unfamiliar conspecific over the familiar one (*shRNA-Scramble*: p < 0.01; *shRNA-Gpr158*: p = 0.62), but not affecting the phenotype of sociability in mice (*shRNA-Scramble*: p < 0.01; *shRNA-Gpr158*: p < 0.01), as shown in Figure 4A-C. Secondly, *rAAV-CamK2A-Cre* was bilaterally injected into the mPFC of *Gpr158^fl/fl^* mice to conditionally knock out *Gpr158* in pyramidal neurons. Of note, *rAAV-CamK2A-Cre* suppressed social preference to the novel conspecific over the familiar one in the social novelty test (*rAAV-CamK2A-Cre*: p < 0.01; *rAAV-Scramble*: p = 0.366), but did not affect their preference to a conspecific over an empty cage in the sociability test (*rAAV-CamK2A-Cre*: p < 0.01; *rAAV-Scramble*: p < 0.05), compared to these mice injected with the virus *rAAV-Scramble* (Figure 4D-F). In both tests, the knockdown of *Gpr158* expression was verified using qPCR assay (Figure 4A and D). These data confirm that GPR158 within pyramidal neurons in the mPFC specifically mediates social novelty, echoing the previous phenotype observed in *Gpr158^-/-^* mice.

**Figure 4.**
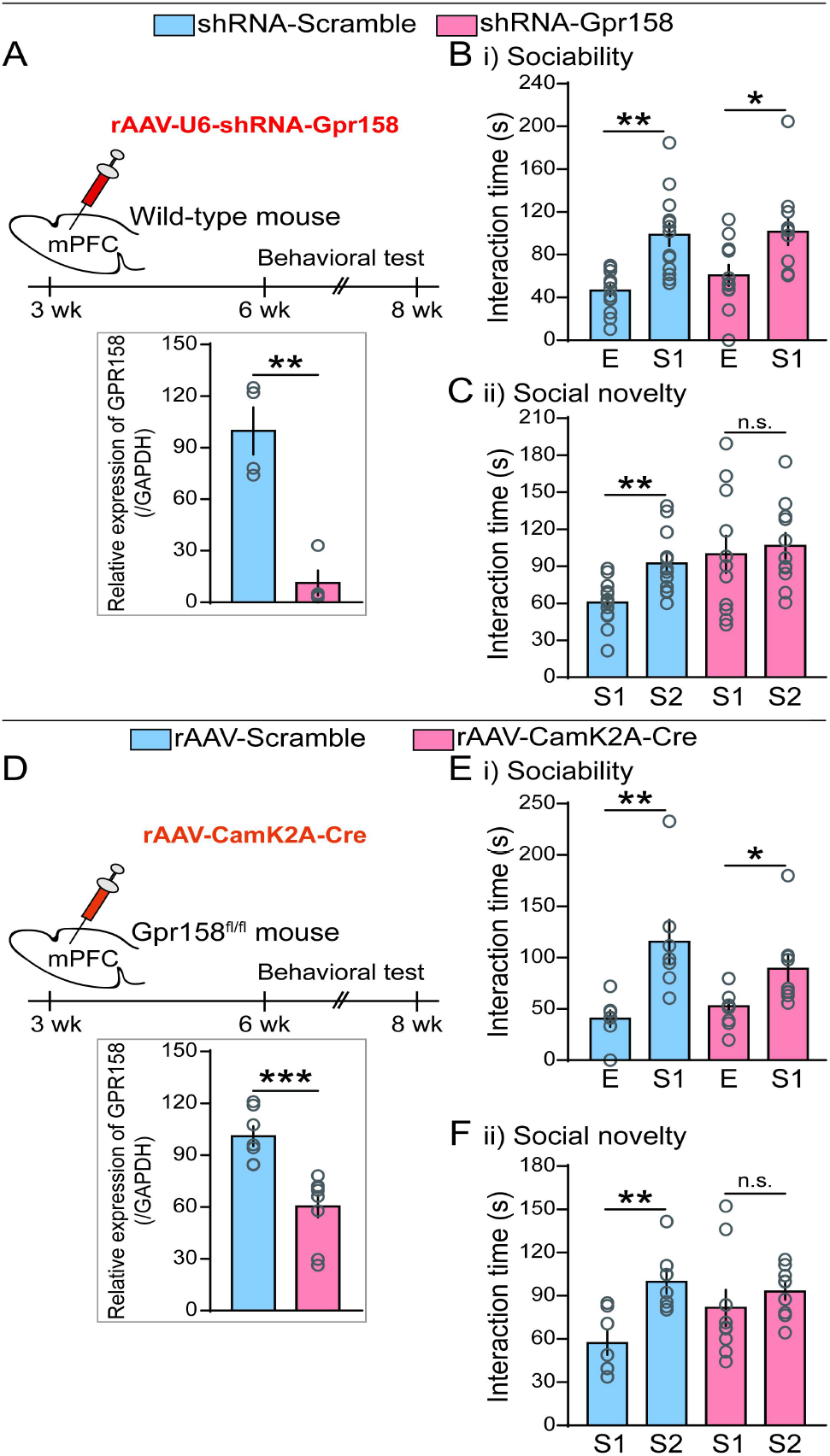
Silencing *Gpr158* expression in the mPFC or deleting *Gpr158* within pyramidal neurons induced social novelty deficit in mice. (A) Knockout of *Gpr158* in the mPFC by *rAAV-U6-shRNA-Gpr158-CMV-EGFP-sv400pA* (*shRNA-Gpr158*) in the WTs compared to those injected with *rAAV-CMV-EGFP-sv400pA (shRNA*-*Scramble)* (*shRNA*-*Scramble*: n=13; *shRNA-Gpr158*: n=11). The knockdown efficiency of *shRNA-Gpr158* was examined by qPCR. Behavioral tests were performed 3 weeks after the virus was bilaterally microinjected into the mPFC (lower panel). (B) Silencing *Gpr158* transcription in the mPFC does not impair sociability behavior (*shRNA-Scramble*: p < 0.01; *shRNA-Gpr158*: p < 0.01). (C) Silencing *Gpr158* transcription in the mPFC induced a phenotype of social novelty deficit in the WTs (*shRNA-Scramble*: p < 0.01; *shRNA-Gpr158*: p = 0.62). (D) Selective deletion of *Gpr158* in pyramidal neurons within the mPFC by *rAAV-CamK2A-Cre* (*rAAV-Scramble*: n=7; *rAAV-CamK2A-Cre*: n=9). The knockout efficiency of *rAAV-CamK2A-Cre* was examined by qPCR. Virus was bilaterally microinjected into the mPFC, while behavioral tests were performed 3 weeks later (lower panel). (E) Conditional knockout of *Gpr158* in pyramidal neurons does not alter sociability behavior (*rAAV-CamK2A-Cre*: p < 0.01; *rAAV-Scramble*: p < 0.05). (F) Conditional knockout of *Gpr158* in pyramidal neurons induced an impaired phenotype of social novelty (*rAAV-CamK2A-Cre*: p < 0.01; *rAAV-Scramble*: p = 0.366). E, an empty cage; S1, stranger 1; S2, stranger 2; F, a familiar object; N, a novel object. Here, n.s., not significant; * p < 0.05, ** p < 0.01. Data were analyzed with One-way ANOVA among 3 groups and the unpaired *t* test between 2 groups. Data are represented as the mean ± SEM.

### Knockout of *Gpr158* disrupted excitatory synaptic transmission in the mPFC

E-I imbalance is the classic mechanism underlying social disorder^2,5^. To assess the potential role of neurotransmitters in modulating the phenotype of social novelty, ELISA was performed to measure the levels of glutamate, GABA and dopamine in the mPFC, hippocampus and serum, respectively (Figure 5A-C). Of significance, the levels of glutamate were significantly decreased in the mPFC (t_10_ = 2.413, p < 0.05) and hippocampus (t_10_ = 2.231, p < 0.05) of KO mice, but unaltered in serum (t_10_ = 2.045, p = 0.067), whereas the levels of GABA were not changed in the mPFC (t_10_ = 1.682, p = 0.123), hippocampus (t_10_ = 0.995, p = 0.343) or serum (t_10_ = 1.256, p = 0.238) of *Gpr158^-/-^*mice. Besides, the levels of dopamine in the mPFC (t_10_ = -2.051, p = 0.67), hippocampus (t_10_ = 0.923, p = 0.378) or serum (t_7.746_ = -1.485, p = 0.177) did not differ between genotypes. Thus, our data indicate that glutamate within the mPFC play a critical role in mediating social novelty.

**Figure 5.**
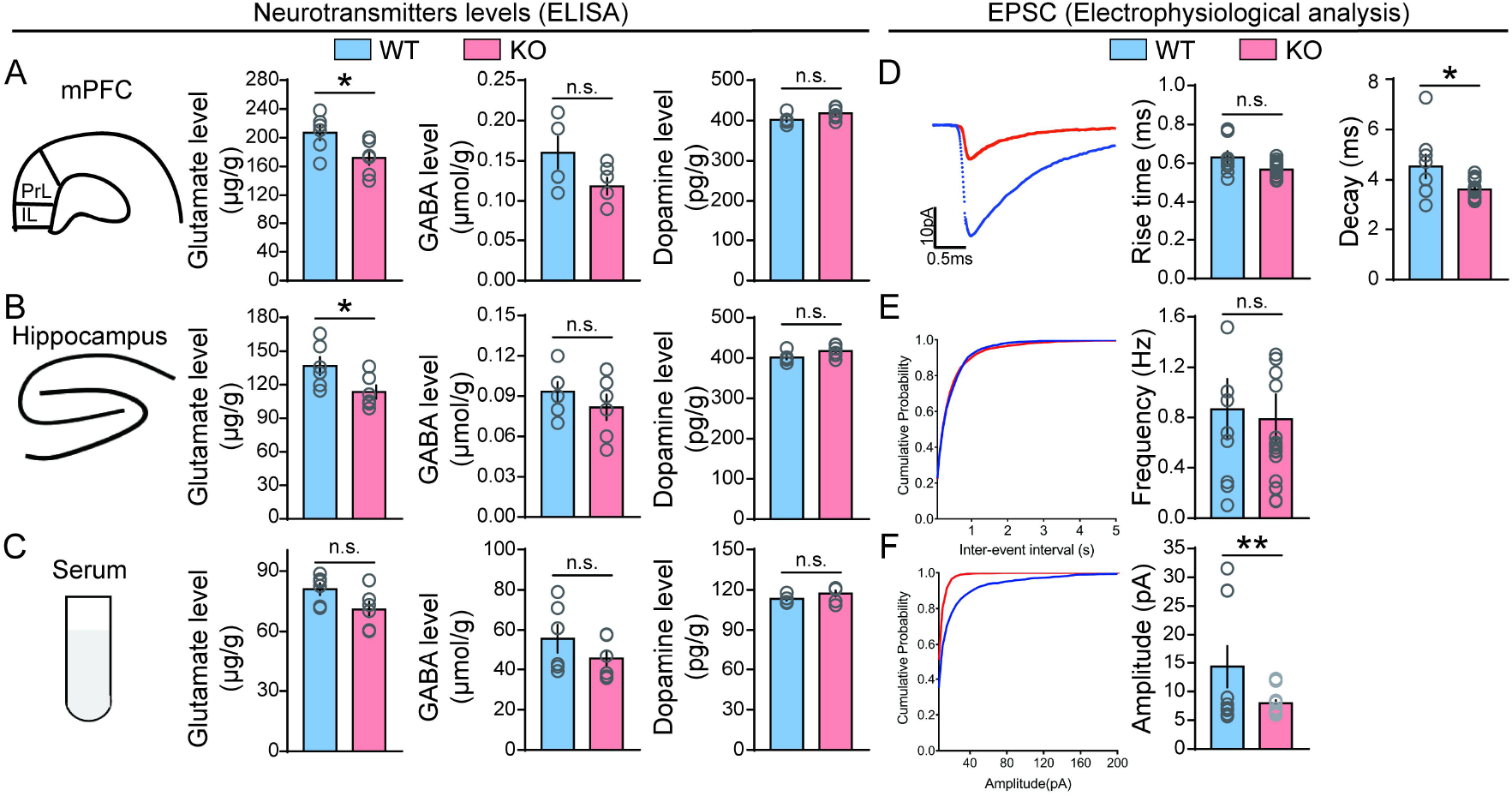
Loss of *Gpr158* interrupted excitatory synaptic transmission in the mPFC. (A) The level of glutamate (t_10_ = 2.413, p < 0.05), instead of GABA (t_10_ = 1.682, p = 0.123) or dopamine (t_10_ = -2.051, p = 0.67), was significantly decreased in the mPFC of KO mice (WT: n=5; KO: n=5). **(B)** The level of glutamate (t_10_ = 2.231, p < 0.05), but not GABA (t_10_ = 0.995, p = 0.343) or dopamine (t_10_ = 0.923, p = 0.378) was decreased in hippocampus of KO mice. **(C)** The level of dopamine (t_10_ = 2.045, p = 0.067), GABA (t_10_ = 1.256, p = 0.238) or dopamine (t_7.746_ = -1.485, p = 0.177) was not altered in serum of KO mice. **(D)** The decay time of EPSCs was significantly decreased in the mPFC of KO mice (t_24_ = 2.931, p < 0.01), while the rise time showed a decreased trend (t24 = 1.948, p = 0.06; WT: n=5; KO: n=5). **(E)** The frequency of EPSCs was not changed (t_24_ = 0.2444, p = 0.809), while **(F)** the amplitude of EPSCs was suppressed in the mPFC of KO mice (t_24_ = 2.339, p < 0.05). WT, wild-type; KO, *Gpr158^-/-^*. n.s., not significant; * p < 0.05, ** p < 0.01. Data were analyzed with the unpaired *t* test. Data are represented as the mean ± SEM.

To validate E-I imbalance in the mPFC, we used whole-cell patch to monitor the neuronal electrophysiological traits^3,21^. In the electrophysiological test, the decay time of sEPSC was found to be shortened in the mPFC of KO mice, while the rise time showed a trend to be decreased compared to that in the controls (Rise: t_24_ = 1.948, p = 0.06; Decay: t_24_ = 2.931, p < 0.01; Figure 5D). The amplitude of sEPSCs was inhibited (t_24_ = 2.339, p < 0.05), while the frequency was not altered in KO mice compared to the WT mice (t_24_ = 0.2444, p = 0.809), shown in Figure 5E and F. The data demonstrate that knockout of *Gpr158* causes functional deficit of excitatory synaptic transmission in the mPFC, potentially accounting for social novelty disorder.

### Knockout of *Gpr158* decreased the density of glutamate vesicles in the mPFC

Rationally, the reduced level of glutamate may be caused by the decreased density of presynaptic glutamate vesicles or glutamatergic synapses^6, 36, 43^. To in-depth reveal whether there were alterations in synapses in the mPFC of KO mice, we observed the synaptic substructures using transmission electron microscopy (TEM). Traditionally, the synapses were briefly classified into two types: asymmetric Gray I synapse (excitatory) and symmetric Gray II synapse (inhibitory) based on their morphology, and then the density of presynaptic vesicles and width of synaptic cleft were measured based on the TEM photographs of synapses^33, 44^. We found that density ofglutamatergic synapses was significantly reduced (t_60_ = 2.203, p < 0.05; Figure 6A), while the number of GABAergic synapses was not altered in the mPFC of KO mice compared to WT mice (t_41_ = 1.913, p = 0.0627; Figure 6B). Meanwhile, there was no difference in width of synaptic cleft of glutamatergic synapses (t_37_ = 0.9125, p = 0.3674) nor in the density of inhibitory vesicles within GABAergic synapses (t_13_ = 1.1039, p = 0.9188) between KO and WT mice (Figure 6A and B). These data suggest that KO mice showed the reduced density of glutamate vesicles in the mPFC, probably accounting for the decreased level of glutamate. To explore whether the density of synapses was reduced, we calculated the density of dendritic spines with Golgi-Cox staining and found no difference in the density of dendritic spines between KO mice and the controls (t_186_ = 1.403, p = 0.1524; Figure 6C). Thus, *Gpr158* knockout could directly result in loss of glutamate vesicles in each synapse, instead of reduce the density of glutamatergic synapses in the mPFC.

**Figure 6.**
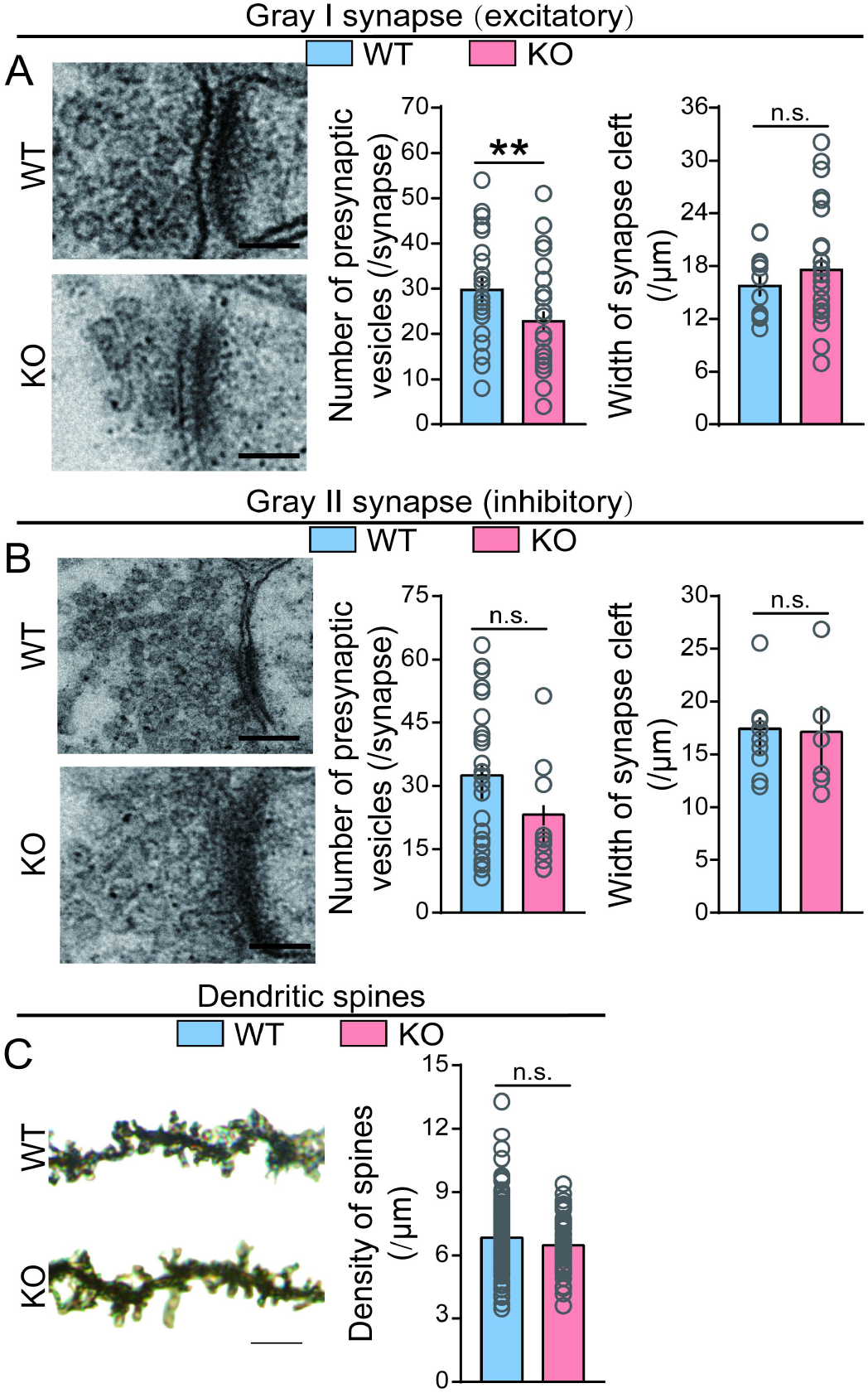
Loss of *Gpr158* decreased the density of glutamate vesicles in the mPFC. (A) Representative images of glutamate vesicles in glutamatergic (Gray I) neurons in the mPFC of KO and WT mice, imaged by TEM. There was a significant decrease in the density of presynaptic vesicles (t_60_ = 2.203, p < 0.05), but not width of synaptic cleft (t_37_ = 0.9125, p = 0.3674) in the mPFC of the KO mice. The images were collected from 4 WT and 4 KO mice. **(B)** Representative images of neurotransmitter vesicles in inhibitory (Gray II) neurons in the mPFC of KOs and WTs, imaged by TEM. There were no alterations in the density of presynaptic vesicles (t_41_ = 1.913, p = 0.0627) or the width of synaptic cleft (t_13_ = 1.1039, p = 0.9188) in the mPFC. Scale bars in **(A)** and **(B)**: 200 µm. **(C)** The density of dendritic spines was not changed in the mPFC by Golgi staining (t_186_ = 1.403, p = 0.1524). Representative images were collected from 4 WT and 4 KO mice. Magnification power 100×. WT, wild-type; KO, *Gpr158^-/-^*; TEM, transmission electron microscopy; n.s., not significant; ** p < 0.01. Data were analyzed with the unpaired *t* test. Data are represented as the mean ± SEM.

### Knockout of *Gpr158* reduces expression of Vglut1 in the mPFC

The inhibited sEPSC amplitude in KO mice might be caused by reduced expression of Vglut1 as a key glutamate transporter located in the vesicular membranes^45, 46^. To explore how GPR158 modulated the presynaptic vesicles, we performed IF and WB tests in the mPFC tissues harvested from KO mice and the controls, and found that Vglut1 expression was significantly reduced (IF: t_233_ = 6.786, p < 0.01;WB: t_7_ = 5.502, p < 0.01), while expression of Gad65/67 was not altered in the mPFC of KO mice (IF: t_213_ = 0.4996, p = 0.6179; WB: t_7_ = 1.009, p = 0.3467), shown in Figure 7. Therefore, we speculate the decreased expression of Vglut1 leads to glutamatergic synaptic transmission in the mPFC.

**Figure 7.**
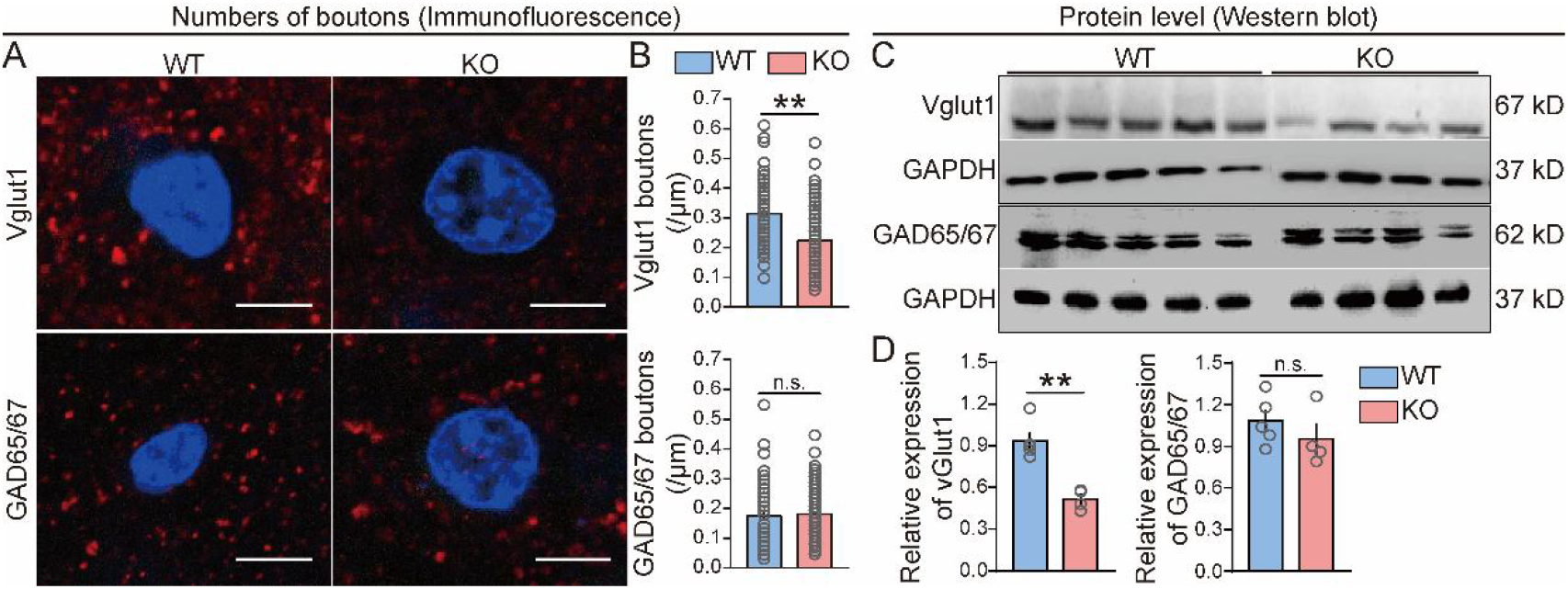
Loss of *Gpr158* inhibited expression of Vglut1 in the mPFC. (A) Representative images of Vglut1 and GAD65/67 boutons in the mPFC of KO and WT mice by immunofluorescent staining (WT: n=5; KO=4). **(B)** In the immunofluorescent staining test, reduced expression of Vglut1 (t_233_ = 6.786, p < 0.01), instead of GAD65/67 proteins (t_213_ = 0.4996, p = 0.6179), was found in the mPFC of KO mice. The density was quantified by the number of boutons distributed in the perimeter. **(C)** Representative images of Vglut1 and GAD65/67 in the mPFC of the KOs and WTs by western blotting assay (WT: n=5; KO: n=4). **(D)** In the western blotting test, the level of Vglut1 (t_7_ = 5.502, p < 0.01), but not GAD65/67 protein (t_7_ = 1.009, p = 0.3467), was decreased in the mPFC of KO mice. Vglut1, vesicular glutamate transporter 1; GAD65/67, glutamic acid decarboxylase at 65 or 67 kD; WT, wild-type; KO, *Gpr158^-/-^*. Scale bars: 5 μm. n.s., not significant; ** p < 0.01. Data were analyzed with the unpaired *t* test. Data are represented as the mean ± SEM.

### Conditional knockout of *Gpr158* in pyramidal neurons led to disrupted expression of V-ATPases and SNAREs in the mPFC

V- ATPases located in the vesicular membrane provide energy to pump glutamate molecules into the synaptic vesicles (SVs), while SNAREs play a role in glutamate fusion and directing glutamate to the presynaptic membrane for exocytosis into the synaptic cleft^17, 45, 47^. To uncover the mechanism underlying deficit of SVs, RNA sequencing was performed using the mPFC from *CamK2A-Cre;Gpr158^fl/fl^* mice and the controls. We found that expression of *SNAREs* such as *Snapin, Stx1α* and *Stx4α*, encoding the vesicular membrane proteins, was significantly reduced, while expression of *Synpr, Stx12, Syt4* and *Sv2b* were enhanced in the mPFC of the mice with cell-specific knockout of *Gpr158*. Meanwhile, expression of V-ATPases including *Atp1a1, Atp2b2, Atp1b2, Atp5f1, ATP6v0c, ATP6v1d, ATP5g3, Atp5d, Atp5o* and *Atp5k* was inhibited, while expression of *Atp5j2, Atp5h, Atp5g1, ATP6v1c, Atp2b4, Atp5a1* and *Atp6v1a* was enhanced in the mPFC of *CamK2A-Cre; Gpr158^fl/fl^* mice (Figure 8A and B). However, we did not find an alteration in the expression of *VGLUT1,* suggesting the changed expression of the protein Vglut1 might be attributed to altered post-transcriptional modifications caused by *Gpr158* deletion.

**Figure 8.**
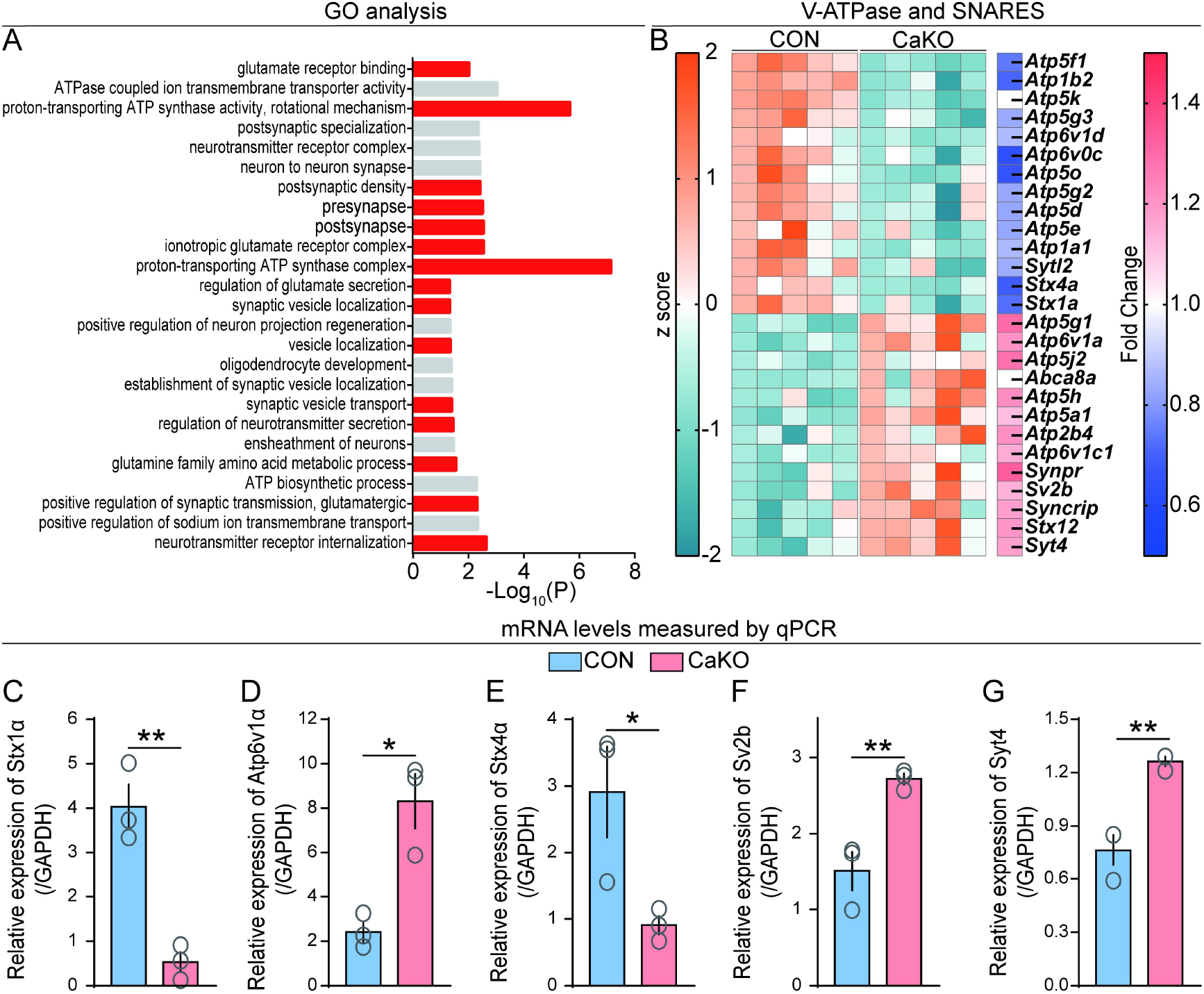
Conditional knockout of *Gpr158* in pyramidal neurons disrupted expression of V-ATPases and SNAREs in the mPFC. GO enrichment and heat map of differentially expressed genes (DEGs) based on RNA-sequencing data from the CaKO mice and the controls (CON). **(A)** Enriched GO pathways. **(B)** Heat map of the DEGs. **(C-G)** Five genes were randomly selected to confirm the expression with the qPCR assay (CON: n=3; CaKO: n=3). Relative expression of *Stx4a, Sv2b, ATP6v1α, Stx1α,* and *Syt4* was altered in the mPFC of CaKO mice (*Stx4α*: t_4_ = 2.885, p < 0.05; *Sv2b*: t_4_ = 4.532, p < 0.05; *ATP6v1α*: t_4_ = 6.295, p < 0.05; *Stx1α*: t_4_ = 4.525, p < 0.05; *Syt4*: t_4_ = 5.514, p < 0.05). GO, Gene Ontology; CON, controls; CaKO, *CamK2A-Cre;Gpr158^fl/fl^*; SNAREs, Soluble N-ethylmaleimide-Sensitive Factor Attachment Protein Receptors. * p < 0.05, ** p < 0.01. Data were analyzed with the unpaired *t* test. Data are represented as the mean ± SEM.

Furthermore, to validate the RNA-Seq findings, 5 genes randomly selected from the DEGs shown in the heat map underwent qPCR tests, and the result was kept consistent *(Stx4α*: t_4_ = 2.885, p < 0.05; *Sv2b*: t_4_ = 4.532, p < 0.05; *ATP6v1α*: t_4_ = 6.295, p < 0.05; *Stx1α*: t_4_ = 4.525, p < 0.05; *Syt4*: t_4_ = 5.514, p < 0.05; Figure 8C-G). In short, GPR158 plays a role in the modulation of the expression of V-ATPases and SNAREs.

## Discussion

GPR158 is known to play a role in synaptic maturation and transmission, which underlies depression and memory loss^3,6,25, 26^. However, its role in social interaction remains unclear. In this study, we demonstrated that GPR158 in pyramidal neurons within the mPFC specifically mediates the phenotype of social novelty. Furthermore, disrupted E-I homeostasis, which is a classic substrate for social interaction, is attributed to the reduced density of glutamate vesicles and suppressed excitatory synaptic transmission within the mPFC of the *Gpr158* knockouts, likely resulted from interrupted expression of Vglut1, V-ATPases, and SNAREs in the membrane of glutamate vesicles^2,11, 48^. Recently, our group have reported that GPR158-positive cells are distributed in many brain areas, particularly cerebral cortex, and only expressed in neurons instead of neuroglial cells, mapped with the transgenic mouse line *Gpr158-e(3×Flag-TeV-HA-T2A-tdTomato)1* which co-expresses epitope-tagged GPR158 and tdTomato^24^. Of interest, most of GPR158 is expressed in CamK2A-positive neurons, while a limited proportion is observed in PV-positive inhibitory neurons, suggesting that GPR158 located in pyramidal or GABAergic neurons might play an essential role in modulating the phenotype of social novelty^24^.

Both enhanced and reduced E/I ratios have been reported to underlie social deficit^49–51^. It has been found that the increased ratio of excitation to inhibition can lead to cortical hyper-excitability and may account for the etiology of ASD with a high comorbidity with epilepsy^42^. On the other hand, the state of hypo-excitation of the PFC and hippocampus can result in social deficit of ASD in the mouse model^52, 53^. For example, *Park* et al found that the significantly reduced E/I ratio underlies sociability deficit in *Cntnap2^-/^*^-^ mice, a mouse model of ASD. Reduced E/I ratio is caused by a relatively decreased level of excitatory neurotransmitter, which may be caused by interruption of neurotransmitter release and re-uptake^46, 49, 52^. Based on our data, we attributed the underpinnings of impaired social novelty to the reduced number of glutamate vesicles at a single-neuron level. Our findings showed that the concentration of glutamate was reduced in the mPFC because of *Gpr158* knockout, while the GABA level was not altered, suggesting that release of presynaptic glutamate vesicles or the density of glutamatergic neurons might be decreased by *Gpr158* knockout at the level of cortical circuit. Additionally, we revealed that the density of pyramidal neurons was not altered in the mPFC of the *Gpr158* knockouts, different from a previously reported study that examined a relatively small number of pyramidal neurons (7 vs 167) and found a decrease in the density of excitatory dendritic spines in the mPFC of *Gpr158* knockouts^21^. Besides, we found that the number of glutamate vesicles was reduced in excitatory synapses within the *Gpr158^-/-^* mPFC. Therefore, we conclude that GPR158 may modulate excitatory synaptic transmission by affecting glutamate vesicles.

Regarding the intracellular mechanism, we observed that the absence of *Gpr158* led to a significant reduction in Vglut1 distribution in the mPFC, as well as disturbance in expression of *SNAREs* and *V-ATPases* in pyramidal neurons^16–19^. Vglut1 plays a role in transporting glutamate, while SNAREs are involved in docking and exocytosis of vesicles into the synaptic cleft^16–19^. V-ATPases provide energy for pumping glutamate molecules into the vesicular space^47^. In short, the altered expression of Vglut1, V-ATPases, and SNAREs collectively affects the density of glutamate vesicles and the extracellular level of glutamate.

GPR158 may modulate the expression of vesicle-related proteins by interacting with RGS7^23^. By using the abovementioned *Gpr158-e(3×Flag-TeV-HA-T2A-tdTomato)1* mice, we confirmed the interaction between GPR158 and RGS7 in mouse brain tissue^25^ (data not shown). As reported, RGS7 is recruited to the plasma membrane by GPR158 and regulates the generation of intracellular AC and cAMP in a stressful state^21, 28^. Thus, GPR158 could modulate intracellular second messenger cAMP and cellular processes by regulating the generation and modifications of vesicle-related proteins through docking to RGS. Recently, glycine has been identified as a ligand for GPR158, and can inhibit the cAMP level by binding GPR158 to modulate cortical neuronal activity^54^. Besides, some studies have shown that the bone-derived hormone osteocalcin (OCN) may modulate GPR158 to alleviate the symptoms of depression, anxiety and memory loss in rodents^55–58^. OCN can activate the brain-derived neurotrophic factor (BDNF) signaling pathway by connecting GPR158 and RbAp48 in the hippocampus^58, 59^.

In conclusion, our results revealed that GPR158 in pyramidal neurons within the mPFC specifically modulates social novelty preference by regulating E-I homeostasis. GPR158 may be a potential target for social disorder treatment.

### Disclosure

The authors declare no conflict of interest.

## Acknowledgements

This work was supported by the grants from the Hundred Talents Program of Sun Yat-sen University (392007, NL), National Natural Science Foundation of China (81874176 and 82072766, NL; 82201698, SW), Shenzhen Sanming Project of Medicine (SZSM201911003, NL), Shenzhen Science, Technology and Innovation Commission (SZSTI) Basic Research Program (JCYJ20190809154411427, NL; JCYJ20210324134800002, SW) and Guangdong Basic and Applied Basic Research Foundation (2021A1515110891, SW). We acknowledge Professor William Richardson, Professor John Wood and Dr. Jing Zhao from University College London for the contribution in correcting this manuscript, and Xiuyan Yang from the Seventh Affiliated Hospital, Sun Yat-Sen University for the contribution to raising mice.

## Author contribution

SW, JC, YL, NL and HL contributed to the experimental design. SW, JC, JJ, JZ and DW performed the experiments. SW, SZ, NL and HL performed data analysis, interpreted data and wrote the manuscript.

## Supplementary Information

**Supplementary Figure 1.**
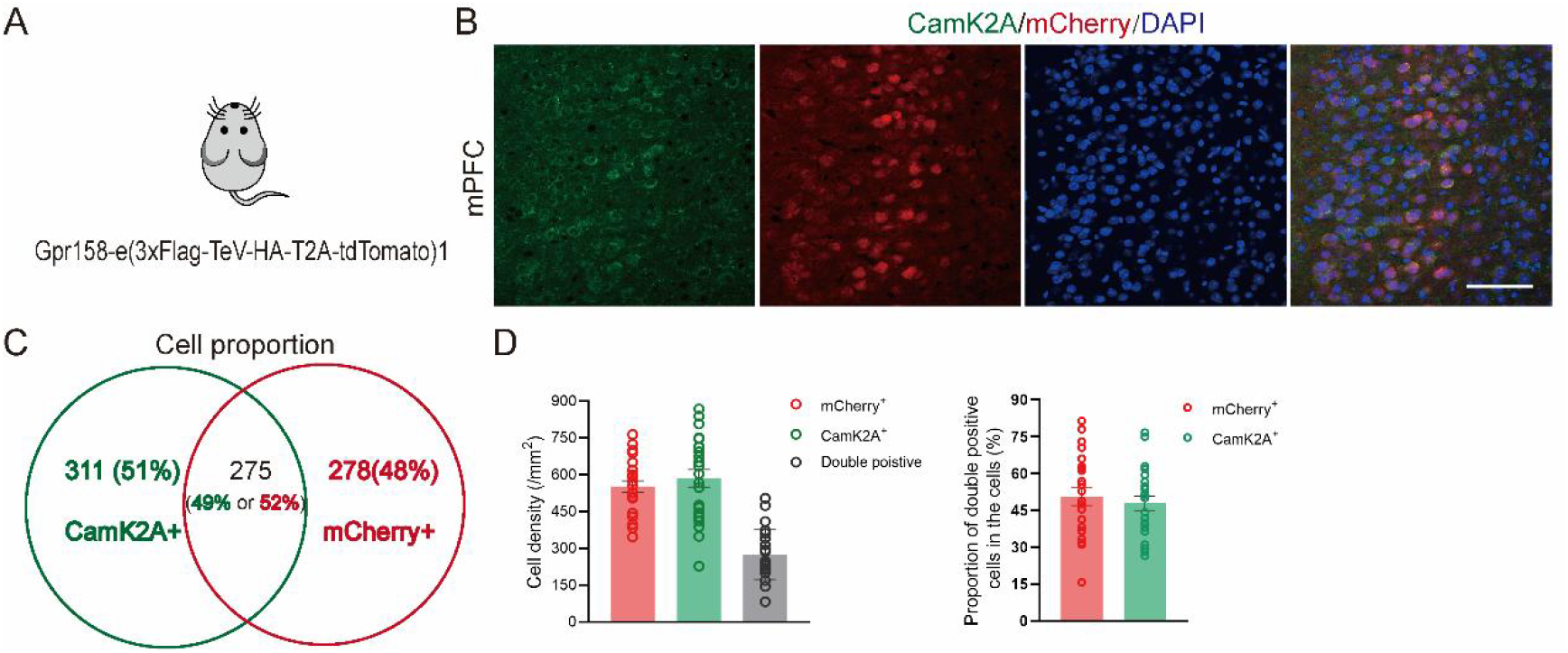
The density of CamK2A^+^ and GPR158^+^ neurons in the mPFC was quantified. (A-B) Representative images of CamK2A^+^ and mCherry^+^ neurons expressed in the mPFC. *Gpr158-Tag* or *Gpr158-e(3×Flag-TeV-HA -T2A-tdTomato)1* mouse was used for immunofluorescent staining. The primary antibody of mCherry targeting tdTomato proteins was used to stain GPR158. **(C)** The proportion of CamK2A^+^ and mCherry^+^ cells. **(D)** Quantification of CamK2A^+^, mCherry^+^ and double-positive cells in the mPFC of *Grp158-Tag* mice (n=4). Scale bars: 20 μm. Data are represented as the mean ± SEM.

**Supplementary Figure 2.**
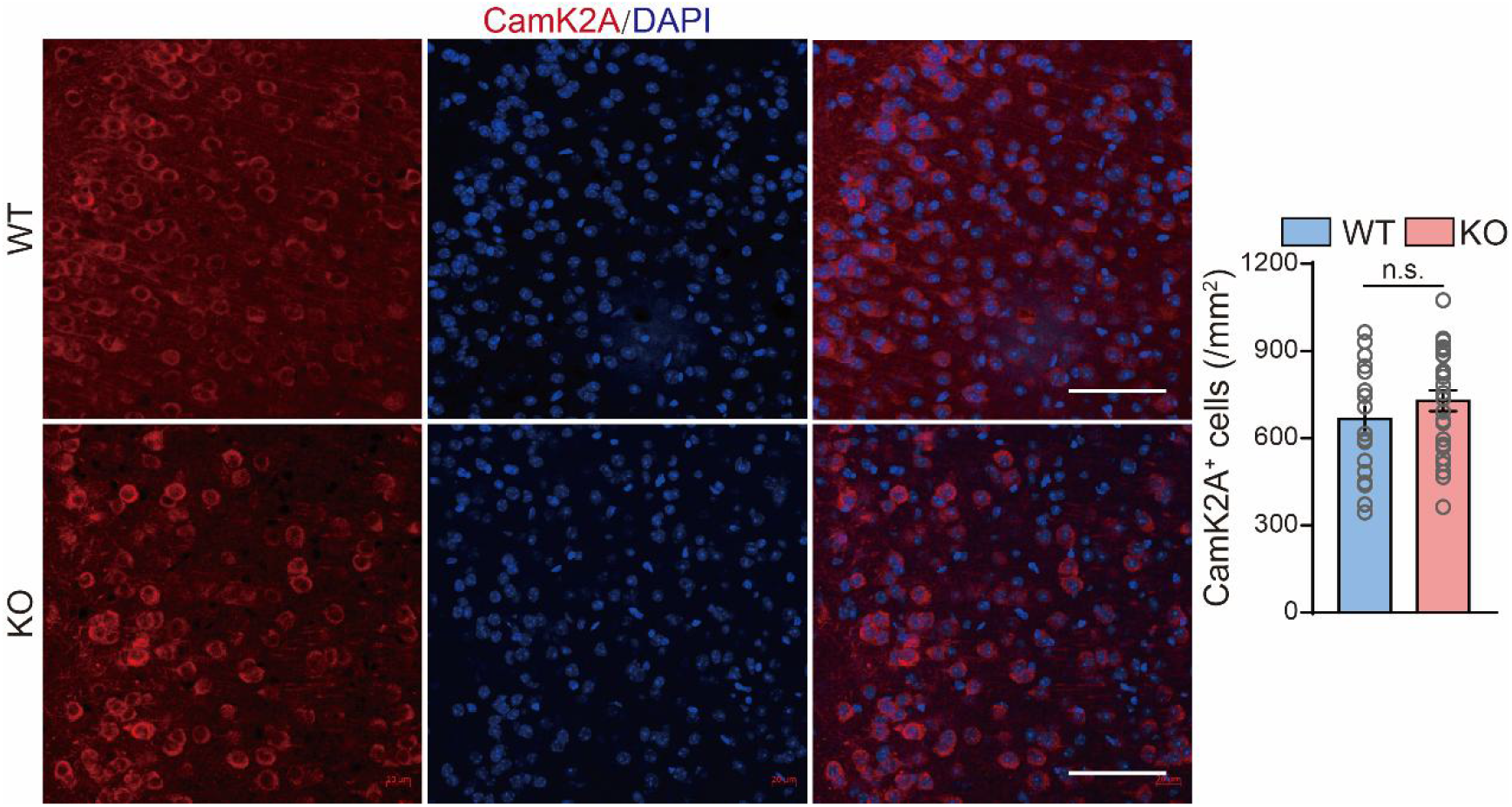
The density of pyramidal neurons was not changed in the mPFC of KO mice. Representative images of pyramidal neurons in the mPFC (left panel). The density of CamK2A^+^ neurons in the mPFC did not differ between KO mice (n=4) and WT littermates (n=4) (t_44_ = 1.125, p = 0.2666; right panel). Scale bars: 20 μm. WT, wild-type; KO, *Gpr158^-/-^*. n.s., not significant. Data were analyzed with the unpaired *t* test. Data are represented as the mean ± SEM.

**Supplementary Table 1.**
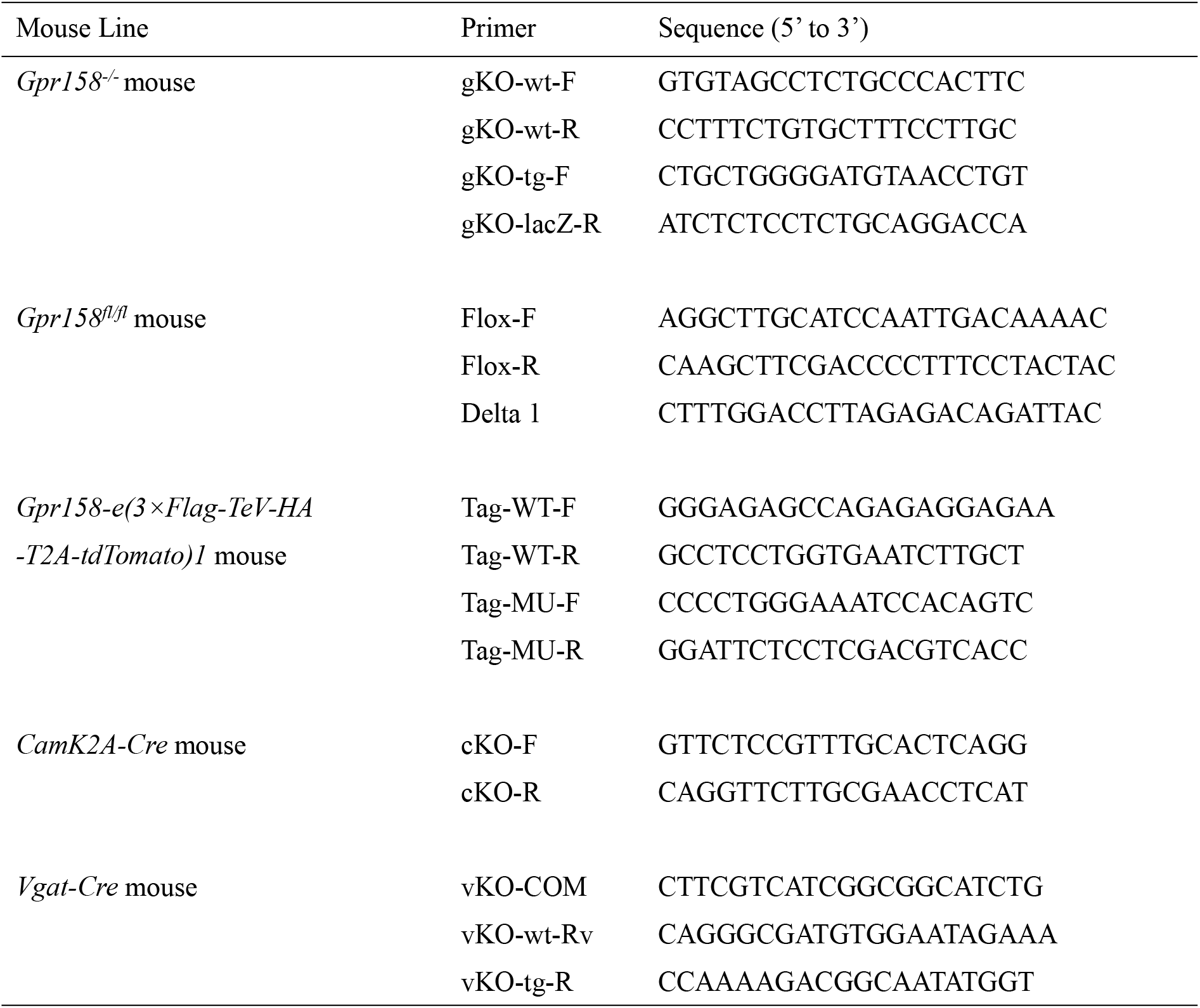
PCR primers for genotyping the genetically modified mice

## Key Resource Table

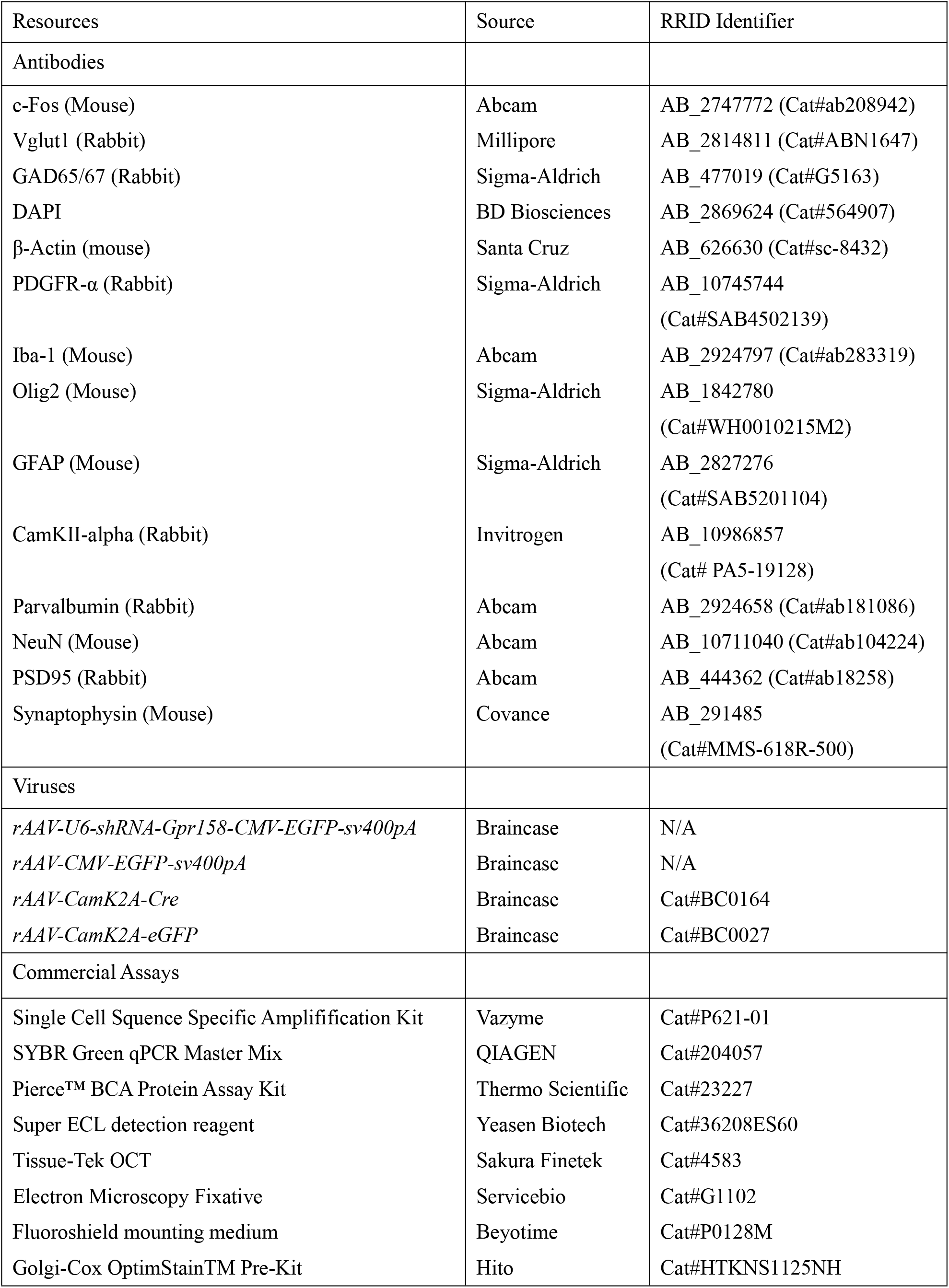

| Resources | Source | RRID Identifier |
| --- | --- | --- |
| Antibodies |  |  |
| c-Fos (Mouse) | Abcam | AB_2747772 (Cat#ab208942) |
| Vglut1 (Rabbit) | Millipore | AB_2814811 (Cat#ABN1647) |
| GAD65/67 (Rabbit) | Sigma-Aldrich | AB_477019 (Cat#G5163) |
| DAPI | BD Biosciences | AB_2869624 (Cat#564907) |
| β-Actin (mouse) | Santa Cruz | AB_626630 (Cat#sc-8432) |
| PDGFR-α (Rabbit) | Sigma-Aldrich | AB_10745744<br>(Cat#SAB4502139) |
| Iba-1 (Mouse) | Abcam | AB_2924797 (Cat#ab283319) |
| Olig2 (Mouse) | Sigma-Aldrich | AB_1842780<br>(Cat#WH0010215M2) |
| GFAP (Mouse) | Sigma-Aldrich | AB_2827276<br>(Cat#SAB5201104) |
| CamKII-alpha (Rabbit) | Invitrogen | AB_10986857<br>(Cat# PA5-19128) |
| Parvalbumin (Rabbit) | Abcam | AB_2924658 (Cat#ab181086) |
| NeuN (Mouse) | Abcam | AB_10711040 (Cat#ab104224) |
| PSD95 (Rabbit) | Abcam | AB_444362 (Cat#ab18258) |
| Synaptophysin (Mouse) | Covance | AB_291485<br>(Cat#MMS-618R-500) |
| Viruses |  |  |
| *rAAV-U6-shRNA-Gpr158-CMV-EGFP-sv400pA* | Braincase | N/A |
| *rAAV-CMV-EGFP-sv400pA* | Braincase | N/A |
| *rAAV-CamK2A-Cre* | Braincase | Cat#BC0164 |
| *rAAV-CamK2A-eGFP* | Braincase | Cat#BC0027 |
| Commercial Assays |  |  |
| Single Cell Sequence Specific Amplification Kit | Vazyme | Cat#P621-01 |
| SYBR Green qPCR Master Mix | QIAGEN | Cat#204057 |
| Pierce™ BCA Protein Assay Kit | Thermo Scientific | Cat#23227 |
| Super ECL detection reagent | Yeasen Biotech | Cat#36208ES60 |
| Tissue-Tek OCT | Sakura Finetek | Cat#4583 |
| Electron Microscopy Fixative | Servicebio | Cat#G1102 |
| Fluoroshield mounting medium | Beyotime | Cat#P0128M |
| Golgi-Cox OptimStain™ Pre-Kit | Hito | Cat#HTKNS1125NH |

